# A Toolbox for Binary and Rheostat-like Modulation of TOX Expression via Genome and Epigenome Editing in Primary Human T cells

**DOI:** 10.64898/2026.08.13.744691

**Authors:** Elaine V. Schanzer, Katharine F. Vostrejs, Alexandra N. Kurciska, Omar Khan, Fyodor D. Urnov

## Abstract

T cells, which are central mediators of the adaptive immune response, can become dysfunctional when faced with persistent antigen stimulation, such as in chronic infections and cancer. This dysfunctional state, known as T cell exhaustion, limits pro-inflammatory T cell function, dampens cytotoxicity and proliferative capacity, and promotes expression of inhibitory receptors. Thymocyte Selection-Associated High Mobility Group Box (TOX) has been proposed as a master regulator of T cell exhaustion due to its necessity for survival of exhausted T cells as well as its role in shaping chromatin accessibility in murine models. Interestingly, partial *Tox* deficiency may improve control of murine tumors. In human tumor infiltrating lymphocytes, high TOX expression is associated with poor disease prognosis. However, the mechanisms by which *TOX* expression is regulated and its importance to human T cell exhaustion remain poorly understood. We report here a robust strategy for generating a genetic knockout of TOX via base editing or a knockout phenocopy via epigenome editing in primary human T cells *ex vivo*, with each approach resulting in near-complete elimination of TOX mRNA. Guided by enhancer prediction data, we use epigenome editing to identify several human *cis*-regulatory regions which function to silence *TOX* expression to varying levels when targeted with CRISPRoff. TOX deficiency had no measurable impact on survival or exhaustion marker levels in human CD8+ T cells in a model of anti-CD3/anti-CD28 stimulation *in vitro*. In agreement with these data, expression profiling revealed that TOX knockout effects on the transcriptome are limited to TOX itself, with no observable downstream effects. These studies show that a complete TOX knockout or silencing has no effect on exhaustion marker expression levels or the transcriptome in repeat-anti-CD3/anti-CD28-stimulated primary human T cells *in vitro*. Taken together, we developed a powerful toolkit of genome and epigenome editing strategies to modify expression of a gene of interest in primary human T cells and study its function. We propose that this framework can be applied to additional genes of interest both to gain mechanistic information about T cell function, as well as develop strategies for improvement of T cell immunotherapies.

## INTRODUCTION

Biologics-based immunotherapies have been transformative in the treatment of a broad range of cancers, with a leading one, pembrolizumab, indicated for 21 different types of cancer, 19 of which are some form of solid tumor and 2 of which are for lymphomas [1]. In striking contrast, the vast majority of approved and in-the-clinic CAR T/TCR T cell therapies are for treatment of relapsed/refractory hematologic malignancies [2]. T cell therapies for solid cancers have struggled to gain traction due to immunosuppression, hypoxia, and competition for nutrients in the tumor microenvironment [2].

The treatment of both solid tumors and hematologic cancers using CAR T approaches face a key biological obstacle: a dysfunctional state known as T cell exhaustion, in which persistent antigen signaling leads to the progressive loss of proliferative capacity, expression of pro-inflammatory cytokines, and cytotoxicity, as well as the upregulation of pro-inhibitory receptors [3–5]. This process is associated with significant epigenetic and transcriptional changes that become increasingly fixed over time and are distinct from other T cell states such as effector or memory states [5–11]. Treatment with immune checkpoint blockade targeting the PD1/PDL1 axis, a strategy that has entered the clinic in combination with CAR T or as a salvage therapy, may temporarily rejuvenate exhausted T cells and induce a more effector-like transcriptional signature, but it fails to epigenetically reprogram exhausted T cells [6,12]. The field could benefit from an enhanced understanding of exhaustion biology as well as strategies to prevent or reverse dysfunction.

In 2019, several groups proposed a central role for the transcription factor TOX (Thymocyte selection-associated high mobility group box), as a master regulator of T cell exhaustion [13–15]. This body of work was inspired by the observation that high TOX expression in tumor infiltrating lymphocytes has been associated with poor prognosis in a number of human cancers [16]. Subsequent efforts largely focused on murine models of T cell exhaustion including chronic infection and tumor models and key results were: 1) TOX was required for the survival of murine T cells in chronic antigen exposure conditions [13,14], 2) TOX was associated with significant epigenetic and transcriptional changes associated with exhaustion in murine T cells [13,14], and 3) partial *Tox* deficiency (*Tox*^+/−^) in murine T cells resulted in improved tumor clearance relative to wild type T cells (*Tox*^+/+^) [13]. To the best of our knowledge the role of TOX in primary human T cell exhaustion has not been investigated directly. Specifically, we are unaware of studies that used the current toolbox of genome and epigenome editing to interfere with TOX expression in primary human T cells and phenotype the consequences.

The present effort was based on the simple premise that, given the formidable body of literature on murine TOX, a phenotype from TOX deficiency in primary human T cells should be straightforward to reveal. To this end we deployed the CRISPR-Cas editing and epi-editing toolkit to modulate *TOX* expression states in primary human T cells and investigate the role of TOX in T cell function during chronic antigen signaling using an established repeat-stimulation model [17,18]. We first established a strategy for near-complete *TOX* knockout in primary human T cells using an adenine base editor. We then established strategies for a comparably near-complete TOX epi-silencing using CRISPRoff by targeting its promoter and also generating a range of *TOX* repression states in rheostat-like fashion by epi-editing *cis*-regulatory elements in the TOX locus. In repeated attempts using T cells from many independent donors, in an established *in vitro* (anti-CD3/anti-CD28) model of repetitive stimulation, *TOX* deficiency had no observable impact on T cell survival, transcriptional signature, or expression of immune checkpoint receptors.

Our studies establish a toolbox for TOX elimination via two orthogonal strategies in primary human T cells, uncover *cis*-regulatory elements of the gene that can be used to establish partial expression epi-states, and show that classical repeat-stimulation of primary human T cells does not reveal any discernible function for TOX for key cell phenotypes.

## RESULTS

We set out to modulate TOX expression in primary human T cells by optimizing editors for genetic knockout (KO) or epi-silencing of TOX. The first approach relied on adenine base editing [19], schematized in Figure 1A (top), which disrupts the endogenous splice donor site (5’-GT-3’) in the first intron of *TOX*, by mutating the A in the second position on the opposite strand to G, resulting in 5’-GC-3’, which is expected to not be recognized by splicing machinery. This is, in turn, expected to result in intron retention, followed by nonsense mediated decay due to the presence of premature stop codons within the retained intron [20,21]. In experiments using T cells from multiple donors, electroporating primary human T cells with the mRNA-encoded ABE8.20m nSpG [22] and splice donor-targeting sgRNA yielded >95% on-target A>G conversion (Fig. S1A-B), resulting in >87% of alleles harboring the “intended edit” splice disruption allele, without any bystanders, as assayed by Next Generation Sequencing (NGS) (Fig. 1B). To interrogate TOX function in primary human T cells via an approach mechanistically orthogonal to base editing we developed a TOX epi-silencing approach, schematized in Figure 1A (bottom). To do so, we turned to CRISPRoff [23,24], composed of catalytically “dead” Cas9 (dCas9) fused to the catalytic domains of DNMT3A, DNMT3L, and KRAB. The DNA methyltransferase domains catalyze CpG methylation, while the KRAB domain recruits KAP1 complex corepressors to form heterochromatin via H3K9me3. When targeted to active gene promoters and other cis-regulatory elements, efficient CpG methylation and H3K9me3 typically result in gene silencing. We delivered CRISPRoff mRNA to primary human T cells along with a pool of 3 *TOX* promoter-proximal sgRNAs selected from the hCRISPRi-v2.1 library [25], “TOX Prom”, to silence *TOX* expression (Fig. 1A), which resulted in robust promoter CpG methylation that spread along the 1.6kb amplicon assayed as measured by enzymatic methylation sequencing (EM-Seq), with several CpG dinucleotides methylated in nearly all reads (Fig. 1C, Fig. S1C-D). By comparison, Mock-electroporated T cells and those which received CRISPRoff and a guide targeting a neutral locus (AAVS1), had little observable CpG methylation at the *TOX* promoter. Both strategies of TOX modulation, genetic KO with ABE and silencing with CRISPRoff, resulted in a robust reduction in *TOX* transcript as measured by reverse transcription quantitative PCR (RT-qPCR) when compared to mock-electroporated controls (Fig. 1D). Taken together our data reduce to practice two orthogonal methods for studying TOX function in human T cells during chronic stimulation.

**Figure 1.**
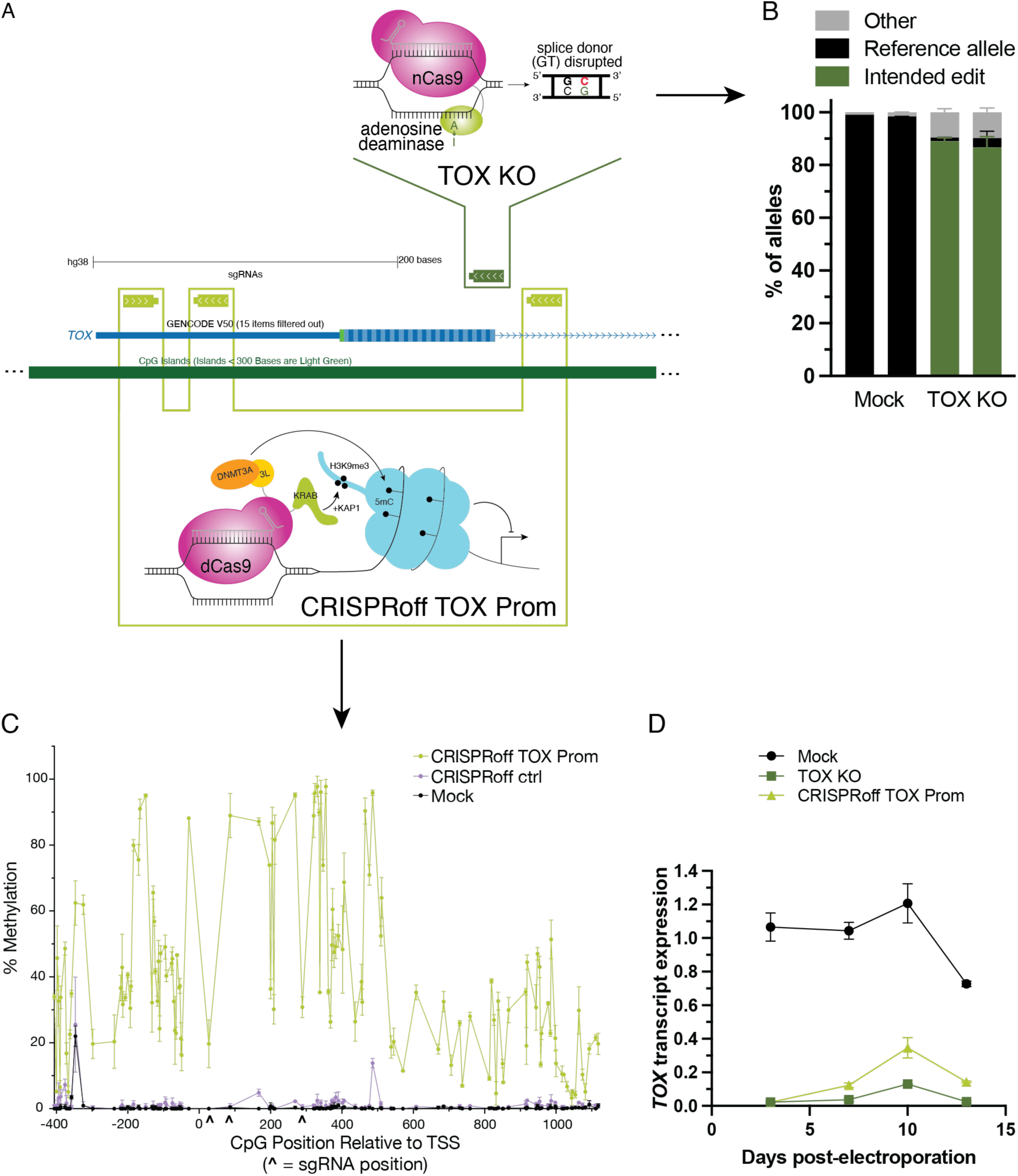
TOX modulation via base editing and epigenetic silencing reduce *TOX* transcript expression in primary human T cells. A) Schematic showing (top) mutation of *TOX* intron 1 splice donor site with ABE8.20m nSpG to generate TOX KO cells; (bottom) targeting of *TOX* promoter with CRISPRoff to induce silencing. B) Quantification of adenine base editing by NGS at intron 1 splice donor. “Mock” refers to mock-electroporated control, TOX KO refers to cells electroporated with ABE8.20m nSpG and an sgRNA targeting the intron 1 splice donor. “Reference allele” refers to the unedited reference sequence, “Intended edit” refers to alleles bearing only the on target A>G edit, and “Other” refers to any alleles bearing bystander edits. C) Quantification of CpG methylation proximal to *TOX* TSS as measured by EM-Seq 7 days after CRISPRoff treatment; N=1 representative T cell donor, with 2 electroporation replicates per treatment condition. D) Representative transcript expression as quantified by probe-based RT-qPCR following ABE and CRISPRoff treatment targeting *TOX.* Transcript expression was calculated by relative quantification with standard curve and normalized to expression of *B2M* transcript. N=1 T cell donor, with 2 electroporation replicates per treatment condition. Each point plotted is an average of at least 3 technical replicates per electroporation replicate.

Complete loss of *TOX* in murine T cells is deleterious to their survival in multiple *in vivo* chronic antigen exposure models, whereas partial loss of *TOX* may be beneficial for tumor clearance [13,14]. While we showed that, in accordance with expectation, epi-editing of the TOX gene promoter yielded near-complete gene silencing, we hypothesized that epi-editing *cis*-regulatory modules of TOX beyond the core promoter would allow us to tune gene expression in a rheostat-like manner to accomplish partial *TOX* knockdown, reasoning that this may be beneficial for T cell survival, while mitigating exhaustion. To do so, we targeted candidate *cis*-regulatory elements (cREs) in the TOX locus for epi-editing, where published evidence for other genes shows that such an approach can potently or moderately repress expression of the corresponding gene [26]. An additional justification for this effort was the hope to gain mechanistic information about regulation of *TOX* expression by probing its candidate *cis*-regulatory elements. To identify such candidate cREs, we mined data from an existing enhancer-gene prediction model, ENCODE-RE2G, and compiled the cREs from several T cell datasets [27], depicted in Figure 2A, bottom track. We targeted CD8+ T cell ENCODE-project-mapped DNaseI Hypersensitive Sites (DHS) that fell within the cREs predicted by ENCODE-RE2G, modules indicated by c01-07 (Fig. 2B) and c08-11 (Fig. 2C), for epi-silencing. We selected a minimum of 4 sgRNAs (and a maximum of 7 sgRNAs) per DHS (Table S1; Fig S2). As shown in Figure 2D, we individually interrogated the contribution, if any, that each cRE makes to TOX gene expression by delivering CRISPRoff mRNA and the sgRNA pools targeting each module to CD8+ T cells (N=2 donors), and then measured *TOX* transcript expression 3 and 7 days (d3 and d7) post-electroporation by RT-qPCR (Fig. 2E and F, respectively). As in the case of a base editing-based TOX knockout (4-5% of control at d3, 7-14% of control at d7), we found that epi-editing TOX proximal to its core promoter resulted in a robust reduction of TOX transcript (3-5% of control at d3, Fig. 2E; 14-19% of control at d7, Fig. 2F). We observed a comparably robust reduction in *TOX* transcript at both d3 and d7 (2-3% of control and 15-23% of control, respectively) by targeting the c04 cRE, which overlaps the region targeted by the Prom pool (Fig. S2C). Whereas c04 guides were distributed across a broad peak in DNase hypersensitivity, the Prom guides (from hCRISPRi-v2.1) are distributed toward the 3’ end of this region, with one guide targeting intron 1 downstream of said peak. In clear contrast, epi-silencing other modules had a markedly non-uniform effect on TOX expression. We observed a marked reduction in TOX transcript when targeting c03, the DHS immediately upstream of the promoter DHS (<0.5 kb upstream of the Transcription Start Site (TSS), Fig. S2C) to 30-33% of control at d3 and 48-58% of control at d7 (Fig. 2E-F). Among the distal modules targeted (>2kb from the TSS), some did not influence TOX expression, while others such as c06 and c10, resulted in moderate TOX repression at d3 and d7 (Fig. 2E-F, respectively; c06: 68-70% and 74-82%, c10 67-82% and 67-72% of control). The sum total of these data demonstrate that, in addition to the core promoter, specific candidate cREs in the TOX locus can be epi-silenced using CRISPRoff with the silenced state maintained over 7 days post-delivery of the mRNA and sgRNAs that form the epi-editor. Further, we show that we were able to, as we set out to do, establish a range of TOX expression states by epi-editing certain, but not all, candidate cREs in the locus.

**Figure 2.**
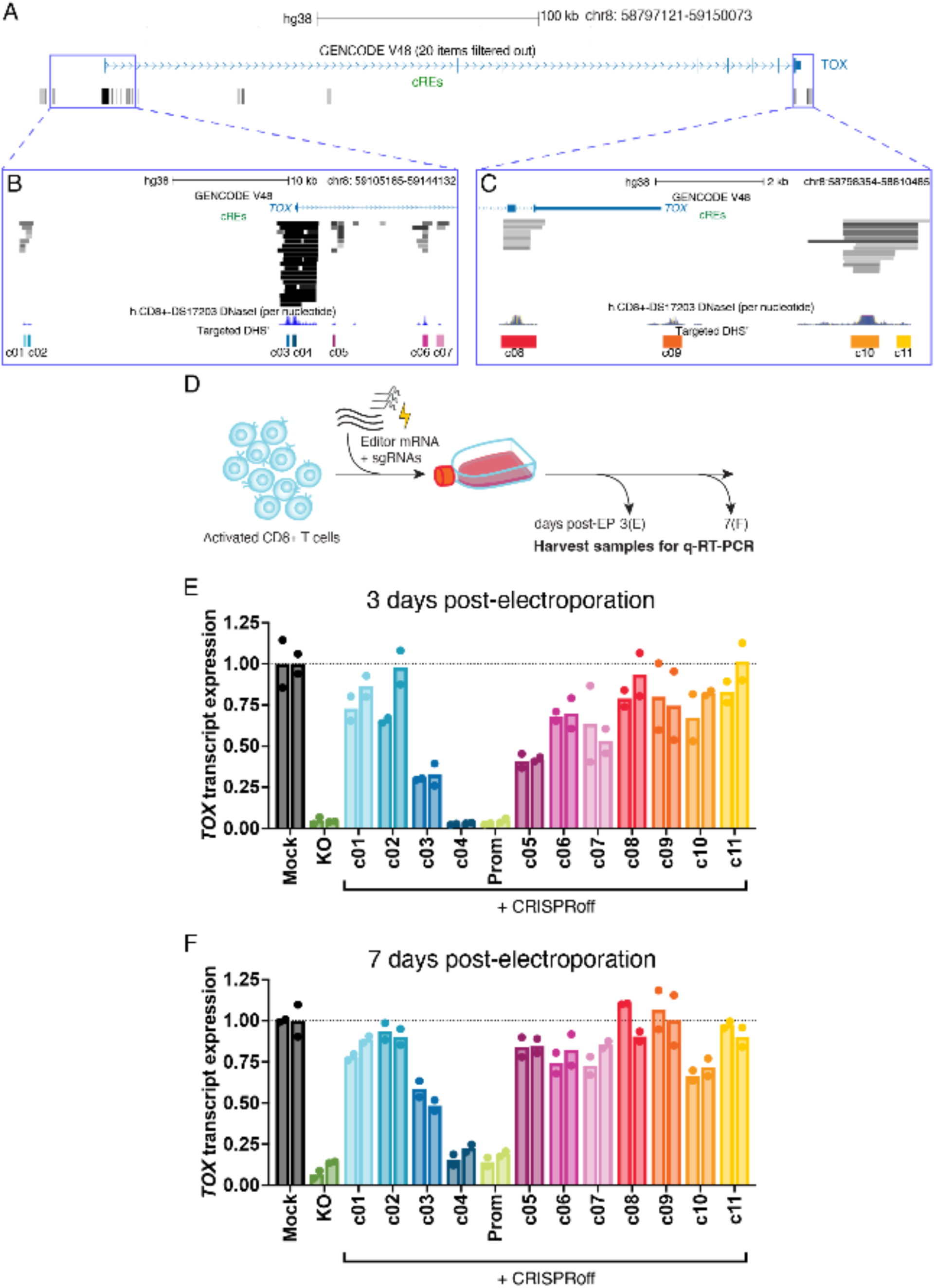
Candidate cis-regulatory elements make unequal contributions to *TOX* expression when targeted with CRISPRoff. (A-C) *TOX* genomic locus visualized in UCSC genome browser regions with track of (A-C) compiled cRE predictions from ENCODE-rE2G model, (B, C) representative DNaseI Hypersensitivity for CD8+ T cells, and (B, C) indicated DNaseI Hypersensitive Sites targeted in study, denoted by c01-11. A) Full *TOX* genomic locus including upstream and downstream accessible regions. B) 5’ distal to intron 1 region, representing targeted DHS’ c01-07. C) Intron 8/Exon 8 to 3’ distal region, representing targeted DHS’ c08-11. (D, E) Transcript expression of *TOX* in primary human T cells transfected with CRISPRoff mRNA and pooled sgRNAs (or indicated controls) at D) 3 days post-electroporation and E) 7 days post-electroporation. “cXX” = CRISPRoff + sgRNA pools targeting individual DHS’ indicated in B and C. Transcript expression was calculated by relative quantification with standard curve and normalized to expression of *B2M* transcript. N=2 T cell donors represented by distinct bars, with 2 electroporation replicates per donor. Each point plotted is an average of at least 3 technical replicates per electroporation replicate. Transcript abundance represented as a percentage of the mock-electroporated control on each day assayed.

Having established 3 complementary modalities of interfering with TOX expression– genetic knockout, complete epi-silencing, and partial epi-silencing– we set out to establish whether partial or complete loss of *TOX* would impact human T cell phenotype during chronic stimulation *in vitro*. We used an established repetitive stimulation model [17,18] in which primary human CD8+ cells were exposed to anti-CD3/anti-CD28 immunomagnetic beads 7, 10, and 12 days post-electroporation (Fig. 3A). We assayed TOX expression 7 days post-electroporation (prior to restimulation) and found, as shown in Figure 3A, a reduction in the percentage of TOX+ cells from 14-21% TOX+ in the mock-treated condition to 1-2% TOX+ cells in TOX KO and TOX Prom treatments. Epi-editing at c06 and c10 reduced the percentage of TOX+ cells to 7-9% and 5-6%, respectively (Fig. S3A). After 3 rounds of restimulation we observed the following: (i) an increase in the percentage of TOX+ cells when comparing mock-electroporated restimulated to non-restimulated (NR) cells (52% and 24% TOX+, respectively); (ii) a reduction in the percentage of TOX+ cells in the TOX KO (20% TOX+) and CRISPRoff TOX Prom (17% TOX+) conditions comparable to non-restimulated condition; and (iii) a moderate reduction in TOX+ cells in cells epi-edited at c06 and c10 cREs (both 34% TOX+) (Fig 3B). Taken together these findings indicate that the expression states imposed by editing and epi-editing (compare to Fig. S3A) are maintained during restimulation. Comparable findings were obtained when this experiment was performed on cells from a distinct donor (Fig S3B). Having comprehensively measured TOX expression across the experiment we then assayed expression of immune checkpoint receptors CD39, LAG3, PD1, and TIM3, following 3 rounds of restimulation. Such markers are upregulated during chronic stimulation in mock-electroporated restimulated cells as compared to non-restimulated (NR) control (Fig. 3B, S3B). We found that neither complete nor partial *TOX* deficiency, however, had a measurable impact on expression of these inhibitory receptors. As shown in Figure 3B (and Fig. S3B), when comparing TOX knockout and knockdown samples (both complete knockdown and partial) to mock-treated control cells, the percentage of CD39+ and LAG3+ CD8+ T cells was largely unchanged; any modest differences in the percentage of PD1+ and TIM3+ cells between treatment groups in one donor were not manifest in cells from a second donor (Fig. 3B, Fig. S3B). PD1 was not strongly induced in either donor tested, making it difficult to draw conclusions about the effect of TOX on its expression (Fig. 3B, Fig. S3B). Taken together, these results suggest *TOX* does not meaningfully influence the expression of canonical exhaustion markers CD39, LAG3, TIM3, and PD1 as measured by flow cytometric immunophenotyping in this specific repetitive stimulation model.

**Figure 3.**
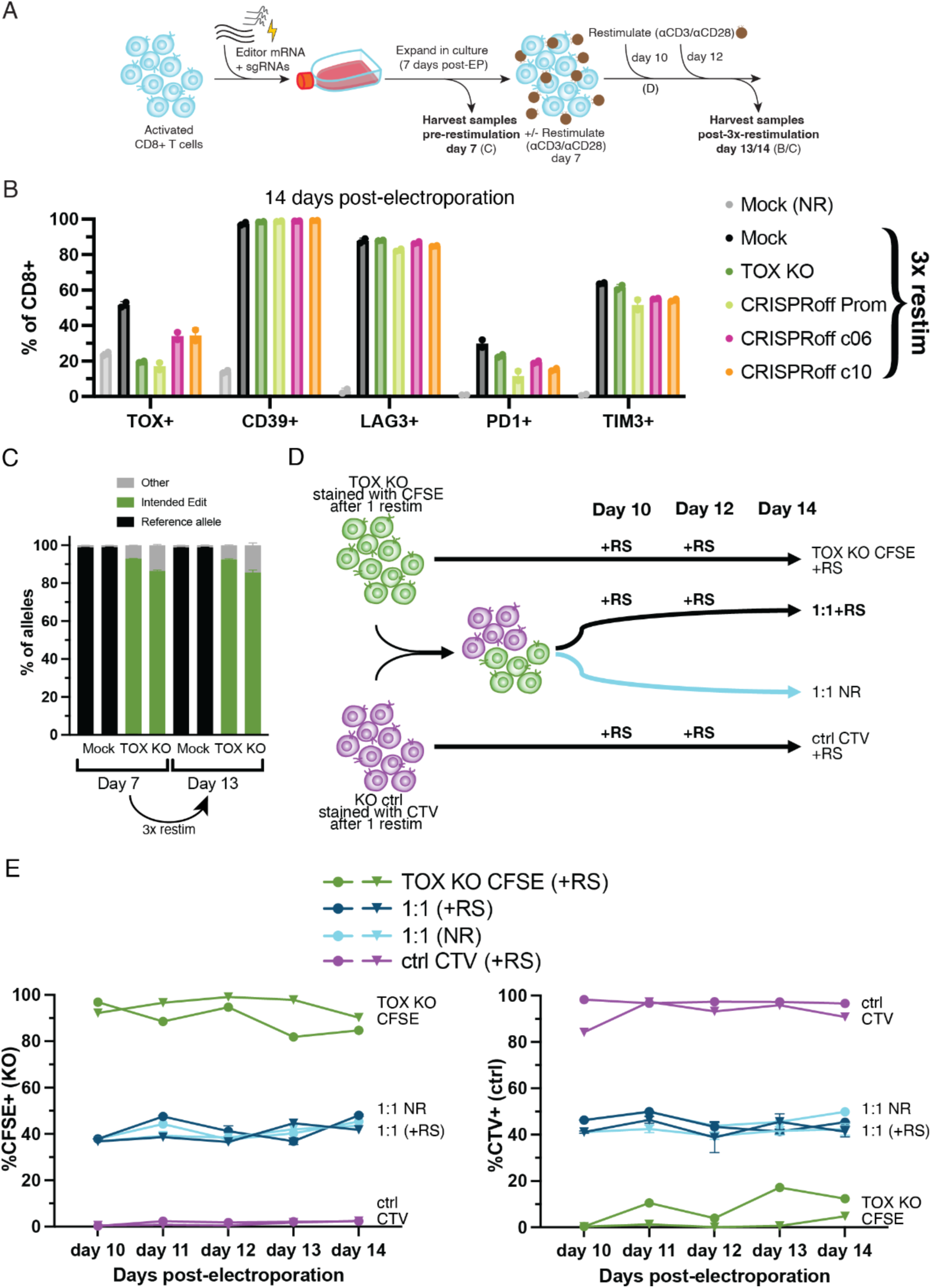
*TOX* deficiency does not result in changes in inhibitory receptor expression nor survival defects in CD8+ T cells repetitively stimulated with anti-CD3/anti-CD28 *in vitro*. A) *In vitro* repetitive stimulation schematic for 3x serial restimulation of edited CD8+ T cells with anti-CD3/anti-CD28 immunomagnetic beads. B) Flow cytometry immunophenotyping of CD8+ T cells from one representative donor 14 days post-electroporation +/− 3x restimulation. Percentage of CD8+ cells expressing CD39, LAG3, PD1, TIM3, and TOX, respectively, were quantified. Bars represent the average of 2 electroporation replicates. C) Tracking of “Reference allele”, “Intended edit,” and “Other” alleles by NGS in unedited (Mock) and TOX KO cells before and after 3x restimulation. D) Schematic depicting co-culture experiment in which TOX KO cells stained with CFSE were mixed 1:1 with negative control (“ctrl”) cells stained with CTV and monitored over time with and without restimulation (+RS and NR, respectively). E) Percentage of CFSE+ (KO, left) and CTV+ (ctrl, right) cells over time as assayed by flow cytometry. “1:1 NR” refers to the 1:1 mix of TOX KO:ctrl cells cultured without further restimulation, “1:1 RS” refers to the 1:1 mix of TOX KO:ctrl cells cultured with restimulation. Each symbol represents cells from a distinct T cell donor (N=2).

Driven by the observation that *TOX* is required for survival of murine T cells in chronic stimulation contexts [13,14], we asked whether *TOX* deficiency impacts human T cell survival in the anti-CD3/anti-CD28 repetitive stimulation model. To do so, we first tracked the frequency of the knockout allele generated by base editing before and after 3x restimulation, as in Fig. 3A (day 7 and day 13, respectively; Fig 3C). We found that the *TOX* knockout allele was maintained both prior to and after 3x restimulation, suggesting no measurable survival disadvantage for *TOX*-deficient cells during chronic stimulation. To explore this issue further, we next asked whether *TOX*-deficient cells would show survival or proliferative defects in direct competition with control edited cells. To do so, we followed a similar repetitive stimulation format as described previously, but on day 10 we stained *TOX* KO cells with CFSE (fluorescent cell tracking dye) and stained control cells (targeted with negative control guide) with CTV (a distinct fluorescent cell tracking dye; “ctrl CTV”) and mixed them together at a 1:1 ratio to track their proliferation and survival over time (as schematized in Fig. 3D). When mixed together, TOX KO CFSE+ cells and ctrl CTV+ cells were each maintained at approximately half of the mixed culture throughout the time course, regardless of restimulation (Fig. 3E). This implies that *TOX* loss is not disadvantageous for neither survival nor proliferation in our restimulation assay, even when TOX KO cells are co-cultured with WT cells. Taken together, our data show that *TOX* deficiency does not appear to influence T cell phenotype with respect to immune checkpoint expression, survival, or proliferation in our hands using an established *in vitro* repetitive stimulation model.

Given that we could not observe any TOX-dependent effect on exhaustion marker expression or cell survival in our assay we reasoned that perhaps such an effect could be observed by genome-wide expression profiling. While the chromatin-wide profile of TOX binding has not been established in human T cells, it is an HMG-box-containing DNA binding protein, the loss of which should be expected to perturb the T cell transcriptome, and in fact does so in murine T cells [13,14]. With a focus on cells with the most robust TOX mRNA depletion via base editing or promoter epi-editing, we performed RNA-Seq on cells with and without *TOX* prior to and following 3x restimulation with anti-CD3/anti-CD28 (Fig. 4A). We included a non-restimulated control cells (Mock (NR)) to account for transcriptional changes over time in culture as well as cells treated with negative control guide to account for the presence of the ABE and CRISPRoff (“KO ctrl”, “CRISPRoff ctrl”, respectively). We first confirmed that *TOX* protein expression remained depleted through day 14 in non-restimulated cells, and in restimulated *TOX* KO and CRISPRoff Prom cells relative to restimulated Mock, KO ctrl, and CRISPRoff ctrl cells via intracellular flow cytometry (Fig. 4B). These expression patterns corresponded with maintenance of robust promoter CpG methylation in the CRISPRoff Prom cells relative to Mock and CRISPRoff ctrl cells (Fig. S4). However, we observed that despite differences in TOX protein expression between day 14 non-restimulated and restimulated mock-treated cells, both populations remained depleted of CpG methylation, suggesting a primed, but inactive chromatin state in the non-restimulated control.

**Figure 4.**
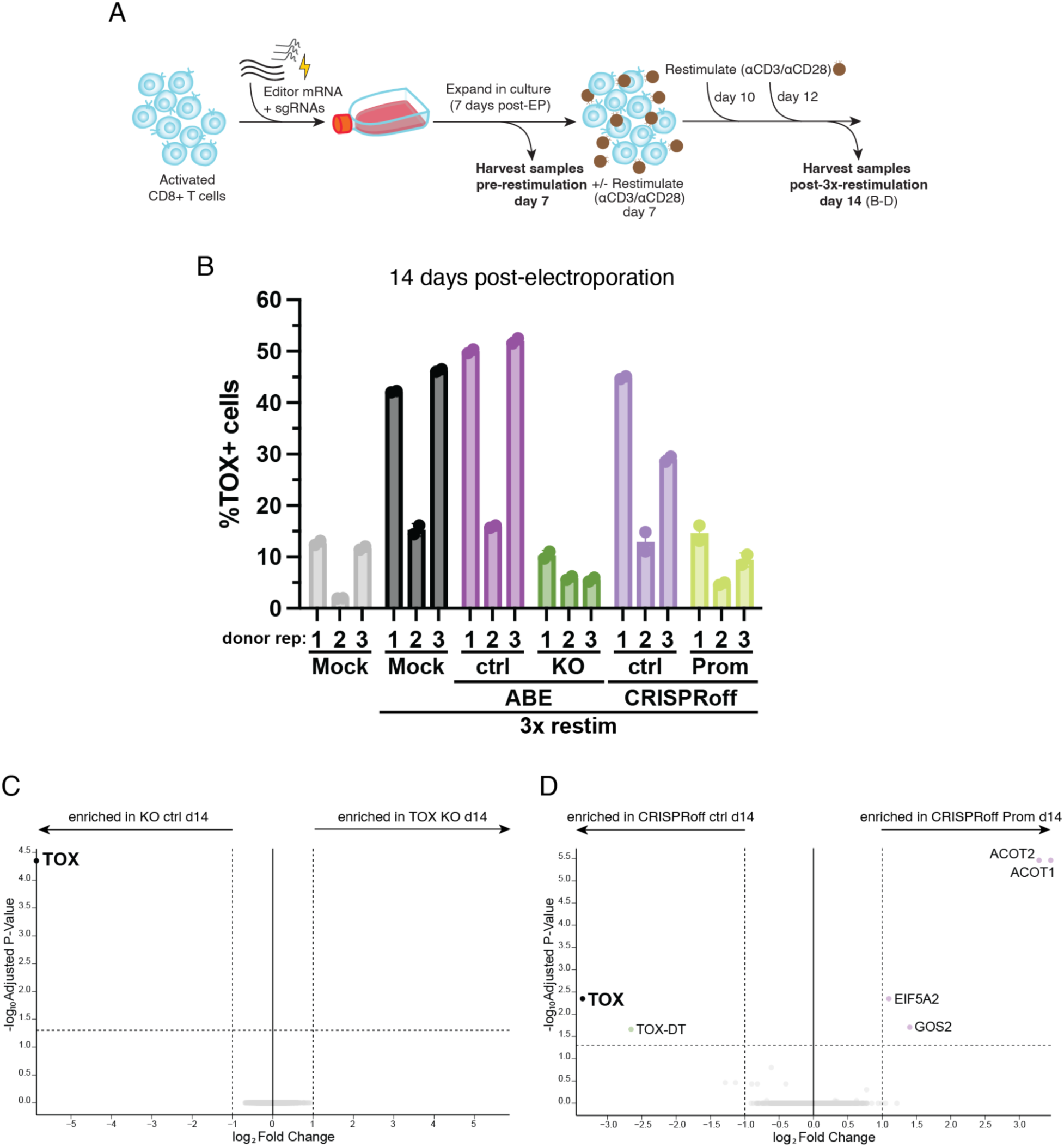
*TOX* deficiency does not impact transcriptional signature of T cells restimulated with anti-CD3/anti-CD28 *in vitro*. A) Experimental schematic; “NR’= non-restimulated. B) Percentage of TOX+ cells 14 days post-electroporation as assayed by intracellular flow cytometry. Each dot represents the average of 2 electroporation replicates from a distinct T cell donor (N=3). C-D) RNA-Seq volcano plots depicting differential gene expression (FC>=2, adj. p-value<0.05) of C) TOX KO and D) CRISPRoff TOX Prom cells relative to their respective negative control; N=3 T cell donors, 2 electroporation replicates.

Analysis of RNA-Seq data showed that, in full agreement with expectation, primary human CD8s experience widespread transcriptional changes during repetitive stimulation (Fig S5). By comparing differential gene expression patterns between mock restimulated and non-restimulated (NR) cells both before and after restimulation, we can observe that in the absence of restimulation, unedited cells downregulate more transcripts than they upregulate (Fig. S5A), whereas upon restimulation, they upregulate many genes between day 7 and day 14 (Fig. S5B), however the starkest contrast in gene expression patterns is between day 14 non-restimulated and restimulated mock cells (Fig. S5C), in which 2224 genes are upregulated and 1803 genes are downregulated. Many transcripts differentially expressed upon restimulation are ones commonly associated with T cell exhaustion signatures, including but not limited to, upregulation of immune checkpoints such as *PDCD1* (PD1), *HAVCR2* (TIM3), *LAG3*, and *ENTPD1* (CD39), and exhaustion-associated transcription factors *TBX21* (Tbet) and *PRDM1* (Blimp1); and downregulation of *TCF7* (Tcf1), associated with T cell stemness. The sum total of this analysis shows that in our assay system primary human T cells respond to restimulation by widespread transcriptional changes affecting genes known to be connected to the biology of T cell exhaustion. We note that *TOX* transcript expression increases modestly over time regardless of restimulation (Fig. S5), and as such it does not appear to be differentially expressed between non-restimulated and restimulated cells at day 14 (Fig. S5C, Mock/Mock NR log_2_FC = 0.27, p = 0.37). This stands in contrast to TOX protein expression, which is indeed upregulated in restimulated cells (35% TOX+) relative to non-restimulated cells (9% TOX+) at day 14 (Fig. 4B). Nonetheless,*TOX* transcript is downregulated in both TOX KO and CRISPRoff TOX Prom cells in comparison to both Mock non-restimulated (“NR”) and Mock restimulated (“Mock”) cells at day 14 (KO/NR log_2_FC = −5.54, p = 1.5 x 10^−11^; KO/Mock log_2_FC = −5.81, p = 7.3 x 10^−9^; Prom/NR log_2_FC = −3.18, p = 2.2 x 10^−8^; Prom/Mock log_2_FC = −3.45, p = 6.2 x 10^−7^).

Having quality-controlled our dataset we asked what transcriptional changes occur when TOX is eliminated by editing or epi-editing. In exceptionally striking contrast to changes driven by restimulation itself (Fig. S4), a comparison of TOX KO cells to KO ctrl cells at day 14 (post-3x-restimulation) identified a single differentially expressed gene: TOX itself (Fig 4C, KO/ctrl log_2_FC = −5.86, p = 3.2 x 10^−9^). This provides independent verification that TOX mRNA is depleted following base editing the *TOX* locus, but clearly indicates that *TOX* knockout does not result in any measurable perturbation to the T cell transcriptome in our assay. As orthogonal confirmation that TOX-deficiency does not perturb the T cell transcriptome, we examined RNA-Seq data from T cells epi-edited at the TOX promoter. This analysis essentially confirmed the findings from TOX knockout cells. Specifically, as shown in Figure 4D, *TOX* is downregulated when comparing day 14 restimulated CRISPRoff Prom cells to CRISPRoff ctrl (Fig. 4D, Prom/ctrl log_2_FC = −3.46, p = 1.3 x 10^−6^). All 4 transcripts upregulated in this comparison are attributable to the presence of the negative control guide, as all 4 genes share a sequence partially homologous to the sgRNA spacer, indicating CRISPRoff repressed these transcripts in a guide-dependent manner in the ctrl-targeted cells but not in the Prom-targeted cells. Further evidence in support of the robustness of our dataset came from the finding that the only other differentially expressed transcript was *TOX-DT*, a divergent lncRNA which is transcribed from the same promoter region in the direction opposite *TOX*. Although the function of this particular transcript is not well-understood, divergent transcripts are generally thought to modulate expression of their neighboring gene, in this case, *TOX*. Because of the proximity of *TOX-DT* to *TOX*, it is not surprising that targeting of CRISPRoff to the TOX promoter would also repress *TOX-DT* (see Fig. S2C). We thus show that repetitive stimulation with anti-CD3/anti-CD28, first established in 1997 [28] and more recently applied to study of T cell exhaustion [17,18], loss of TOX driven by two orthogonal methods fails to perturb the T cell transcriptome, has no effect on T cell survival *ex vivo*, and has no effect on expression of exhaustion markers.

## DISCUSSION

Here we describe several strategies of TOX modulation with CRISPR-Cas and explore their impacts on the function of chronically stimulated primary human CD8+ T cells *in vitro*. We established a base editing-mediated knockout of TOX by mutating its splice donor site in intron 1 (Fig. 1 & Fig. S1A-B). We also established a near-complete knockdown of TOX by targeting CRISPRoff to the promoter region of the TOX locus (Fig. 1 & Fig. S1C-D). Finally, by targeting candidate TOX *cis*-regulatory elements with CRISPRoff, we established a range of intermediate TOX expression epi-states (Fig. 2). We were not able to observe phenotypic differences between TOX-deficient and control cells when assaying for expression of immune checkpoint receptors (Fig. 3, Fig. S3), survival (Fig. 3), nor transcriptional differences (Fig. 4) in an established repetitive stimulation model with anti-CD3/anti-CD28. We did not observe survival defects in completely TOX-deficient (TOX KO) human T cells in our repetitive stimulation model using anti-CD3/anti-CD28, which stands in contrast to the finding that TOX is required for the survival of chronically stimulated murine T cells [13,14]. In line with our findings that TOX deficiency did not impact human T cell survival, TOX-deficient human T cells were not significantly enriched in previous genome-wide CRISPR KO screens that tested for improved proliferation in immunosuppressive conditions [29] nor survival after repetitive tumor stimulation [30]. If human TOX is, in fact, required for the survival of primary human CD8+ T cells, this requirement does not reveal itself in repeat-stimulation assays in our work, nor in CRISPR KO screens.

TOX deficiency did not contribute to perturbations in the transcriptional signature of chronically stimulated T cells with a genetic knockout of TOX nor epi-silencing of TOX via targeting of its promoter in our *in vitro* model (Fig. 4). In murine T cells, loss of TOX was associated with widespread changes to chromatin accessibility and transcript expression in distinct models of chronic stimulation [13,14]. Because we did not observe any functional differences in transcript expression between human T cells with and lacking TOX, we did not pursue characterization of chromatin accessibility in our system. Furthermore, we did not observe differences in immune checkpoint inhibitor expression when investigating cells with TOX KO, complete epi-silencing, nor partial epi-silencing.

Taken together, we were unable to identify a role for TOX in this model of chronic stimulation in human CD8+ T cells using survival, transcriptomic, or exhaustion marker expression. We chose to focus on repetitive stimulation with anti-CD3/anti-CD28, a well-established model, based on findings that T cell exhaustion is TCR-dependent [31]. Thus we selected a reconstitution model intended to focus on T cell-intrinsic epigenetic regulation involving TOX downstream of TCR signaling. More complex models, such as those with antigen-expressing tumor cells, may, however, provide extrinsic cues such as inhibitory ligands or immunosuppressive cytokines that could be important for studying the function of TOX in human T cells [29,30,32]. Our studies provide a comprehensive toolbox for editing and epi-editing human TOX that can be used to create a range of TOX expression states in primary human T cells in any assay system.

Mechanistic knowledge of enhancer biology led to the approval of the first CRISPR gene-edited therapy, Casgevy, which targets the erythroid-specific enhancer of BCL11A with Cas9 nuclease to prevent repression of fetal hemoglobin [33]. As such, modification of *cis*-regulatory elements is therapeutically tractable. Here, instead of using Cas9 nuclease, we take an epi-silencing approach with CRISPRoff. While we did not uncover a TOX-specific phenotype in primary human T cells in our studies, we did begin to unravel regulation of TOX expression by functional interrogation of candidate *cis*-regulatory elements with CRISPRoff (Fig. 2). Such an approach can be deployed on other genes with proposed involvement in T cell function.

Treatment with CRISPRoff leads to targeted CpG methylation (catalyzed by DNMT domains), as we and others have shown, as well as H3K9me3 deposition, a mark of repressive chromatin (from recruitment of chromatin remodelers via the KRAB domain). In our studies, c03-, c04-, and promoter-targeting guide pools overlapped a promoter-proximal CpG island and contributed to robust silencing of TOX expression. When targeting CRISPRoff to the TOX promoter, we observed spreading of CpG methylation along the 1.6kb region assayed, which spanned 0.4kb upstream of the TSS to 1.1kb downstream of the TSS (Fig. 1, Fig. S1C-D, and Fig. S4). CpG methylation resulting from CRISPRoff treatment was maintained over 2 weeks in culture and 3 rounds of restimulation with anti-CD3/anti-CD28 and correlated with TOX silencing that was maintained at both the mRNA and protein level. Given the close proximity of c03 to the TSS (<0.5kb upstream, Fig. S2C) and that repressive chromatin marks from epi-editing can spread [34,35], we are unable to conclude whether the silencing we observed is due to spreading of repressive marks from c03 to the core promoter region, or a distinct *cis*-regulatory mechanism. Targeting of distal elements like c06 and c10, which contain low CpG density, contributed to moderate TOX epi-silencing. Neither of these modules contains a conventional CpG island, pointing to a potential mechanism of long-term epi-repression via a process other than conventional hypermethylation. A simple possibility is that a looping-based contact between a distal cRE and the TOX promoter brings the epi-silencing machinery bound to the former in spatial proximity to the latter; we note that cREs investigated in our studies were nominated using a model that incorporates 3D contact data. More broadly, the extent to which CpG density contributes to epi-editor-mediated-silencing at promoter and distal *cis*-regulatory elements is not well understood, however we note examples of CpG-poor promoters which are strongly and durably silenced by CRISPRoff [24].

At a mechanistic level, there is much left to learn about TOX, its regulation, and its role in human T cell exhaustion. It has been proposed that TOX acts primarily as a repressor of effector gene programs, due to increased accessibility at effector-associated sites in TOX-deficient murine T cells [13]. Alternatively, CUT&RUN assays in murine T cells suggest that TOX binding is largely associated with activation of progenitor and terminal exhaustion gene programs in progenitor and terminally exhausted subsets, respectively [36]. To our knowledge, genome-wide occupancy of TOX has yet to be mapped in primary human T cells. As previously mentioned, exhaustion-specific chromatin accessibility diverges between human and murine T cells, so determining where TOX binds in human T cells may be informative in understanding potential differences between the role of human versus murine TOX in T cell exhaustion, but as previously mentioned, finding appropriate human T cell exhaustion models for studying TOX will be necessary.

We largely focused on the role of TOX in chronically stimulated CD8+ T cells, because they are the cell type canonically associated with exhaustion. However, we note emerging evidence suggests a role for TOX in memory subsets [37] and some CD4+ T cell subsets [38]; with TOX potentially promoting, rather than dampening, anti-tumor activity in CD4+ T_H_1. Whether or not CD4+ T cells experience bona fide exhaustion is not well established [39,40], but the interplay between CD4+ and CD8+ T cells is crucial in many disease processes, and may also be important here.

Furthermore, it has long been appreciated that TOX is required for thymic development of CD4+ T cells in mice [41–44]. TOX is highly conserved between humans and mice (∼90% identical at amino acid and nucleotide level [42]), however, no known inborn errors of immunity in humans are associated with variants in TOX [45,46]. Overall, human genetic variation at the TOX locus is not well understood. No single nucleotide variants in TOX are annotated as Pathogenic/Likely Pathogenic in ClinVar; almost all annotated variants are of uncertain significance and a majority do not specify a related condition [47]. Two published variants in non-coding regions of TOX (one in intron 8, one downstream of the 3’UTR) are associated with pulmonary tuberculosis risk [48]. The mechanism by which these variants influence disease etiology is unknown, but it has been hypothesized that such variants may cause defects in CD4+ T cells, which are crucial for *M. tuberculosis* control.

The tools we optimized for knockout and knockdown of TOX in our studies can be applied to studying the function of TOX in additional exhaustion models or other contexts. We established moderate TOX silencing in particular because partial TOX deficiency (Tox +/−) in murine T cells led to enhanced tumor control [13], implying a dosage-dependency. Despite not observing a phenotype in our system, the role of partial and complete TOX deficiency in chronic stimulation remains an area of open investigation.

---

**Supplementary Figure 1.**
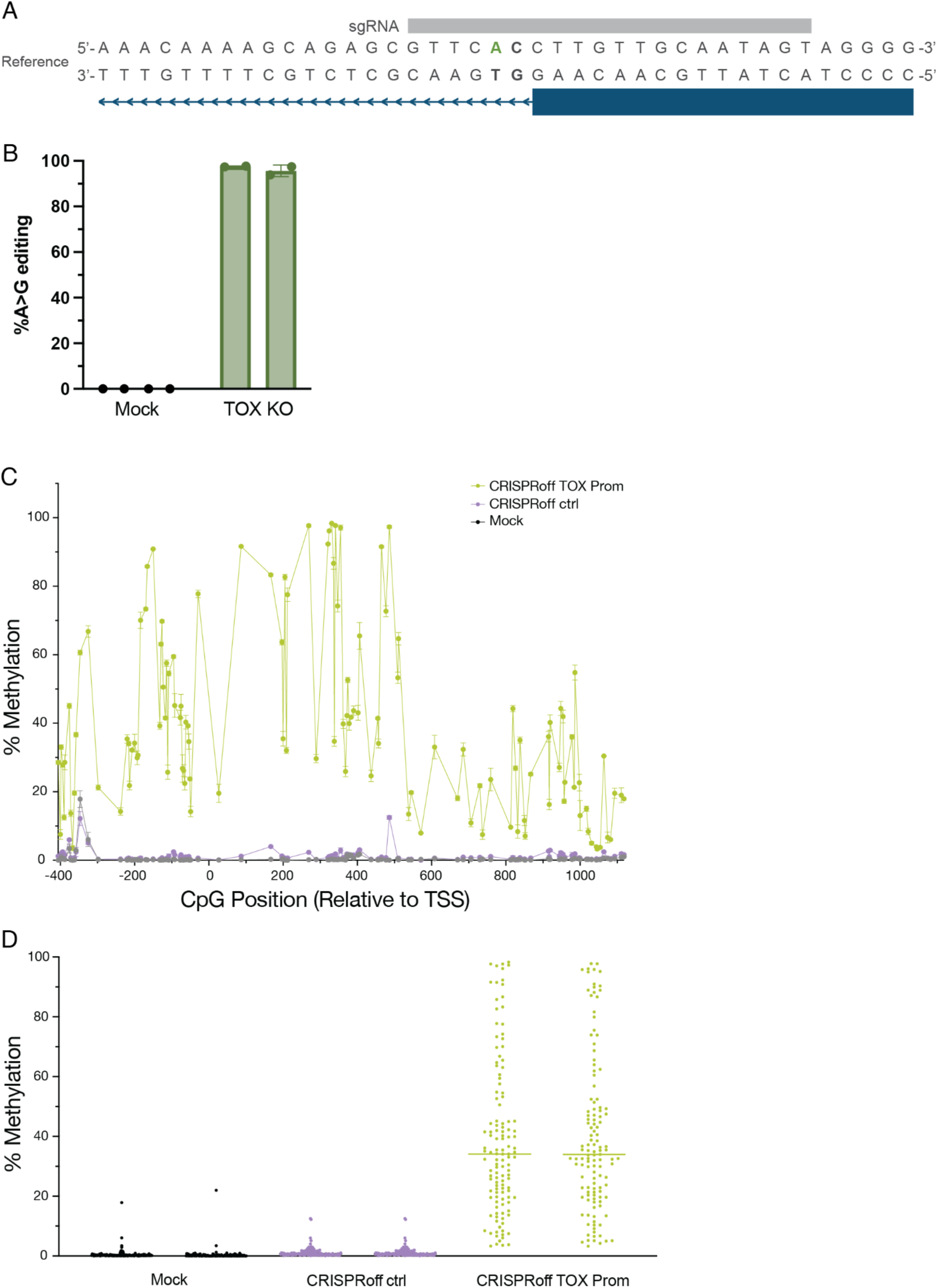
TOX modulation via splice disruption and promoter methylation. A) Schematic for disruption of *TOX* intron 1 splice donor with adenine base editing. B) Quantification of total on-target A>G editing at *TOX* intron 1 splice donor by NGS. C) Quantification of spatially-resolved CpG methylation proximal to *TOX* TSS as measured by EM-Seq 7 days after CRISPRoff treatment; N=1 T cell donor (compare to Fig. 1C), with 2 electroporation replicates per treatment condition. D) Average percent methylation per CpG dinucleotide in *TOX* amplicon as measured by EM-Seq 7 days after CRISPRoff treatment; N=2 T cell donors with 2 electroporation replicates per treatment condition.

**Supplementary Figure 2.**
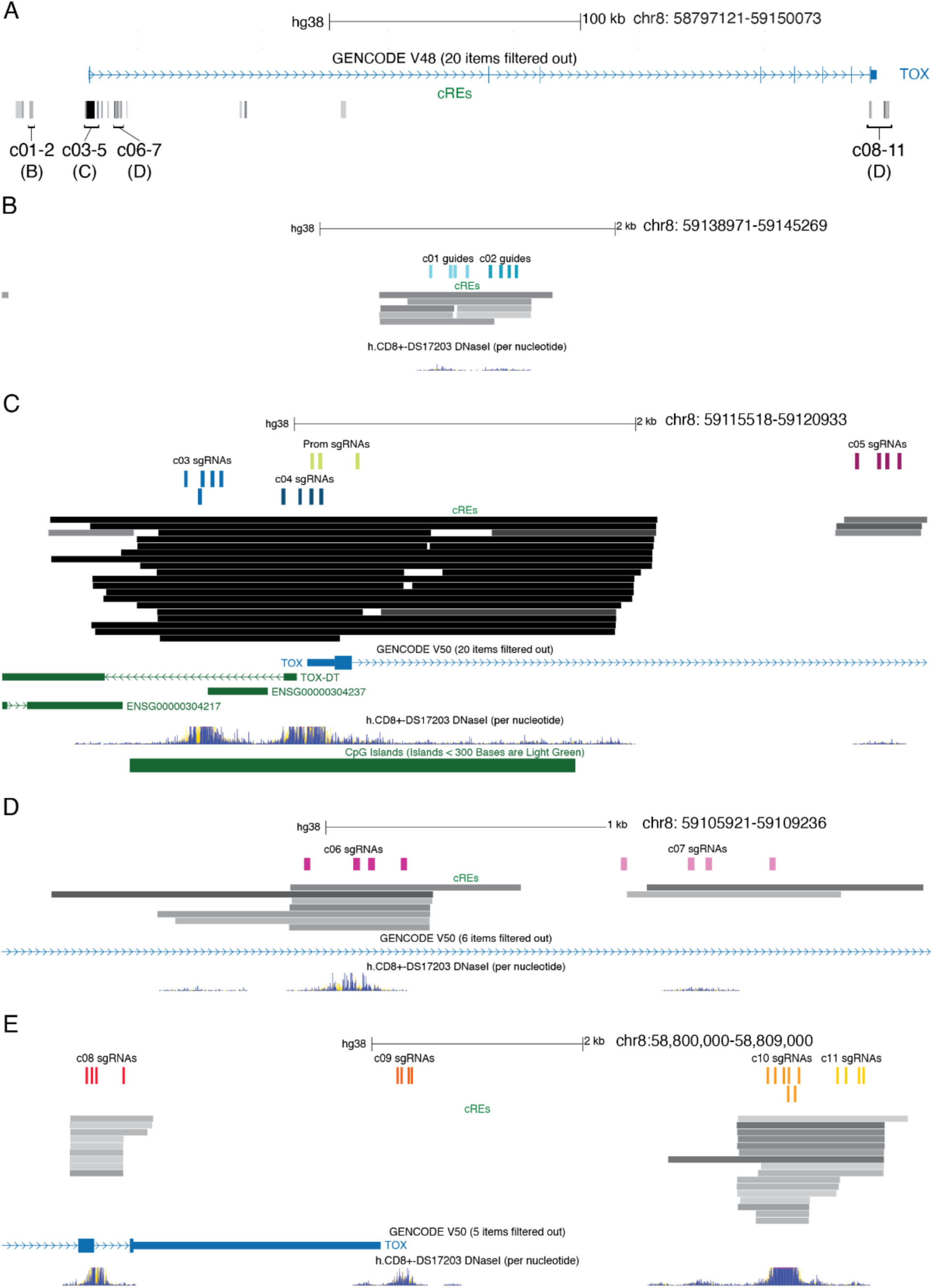
Targeting *TOX* candidate *cis*-regulatory elements with CRISPRoff by tiling guides across individual DNaseI Hypersensitive Sites. A) Schematic of full *TOX* genomic locus and candidate *cis*-regulatory elements (cREs) with 5’ distal (B), TSS proximal (C), Intron 1 (D), and 3’ distal (E) regions indicated. B) 5’ distal cREs c01 and c02 with sgRNA target sequences indicated. C) TSS proximal region with c03, c04, Prom, and c05 guides indicated. “Prom” guides were sourced from hCRISPRi-v2.1 library and overlap with guides in c04, both groups target the DHS most proximal to the *TOX* TSS. The DHS targeted by c05 guides is located within intron 1. D) Intron 1 cREs c06 and c07 with guides indicated. E) 3’ distal region with c08 targeting a DHS overlapping TOX exon 8/intron 8, c09 targeting a DHS at the end of the 3’UTR (not overlapping predicted cRE), c10 and c11 targeting 3’ distal DHS.

**Supplementary Figure 3.**
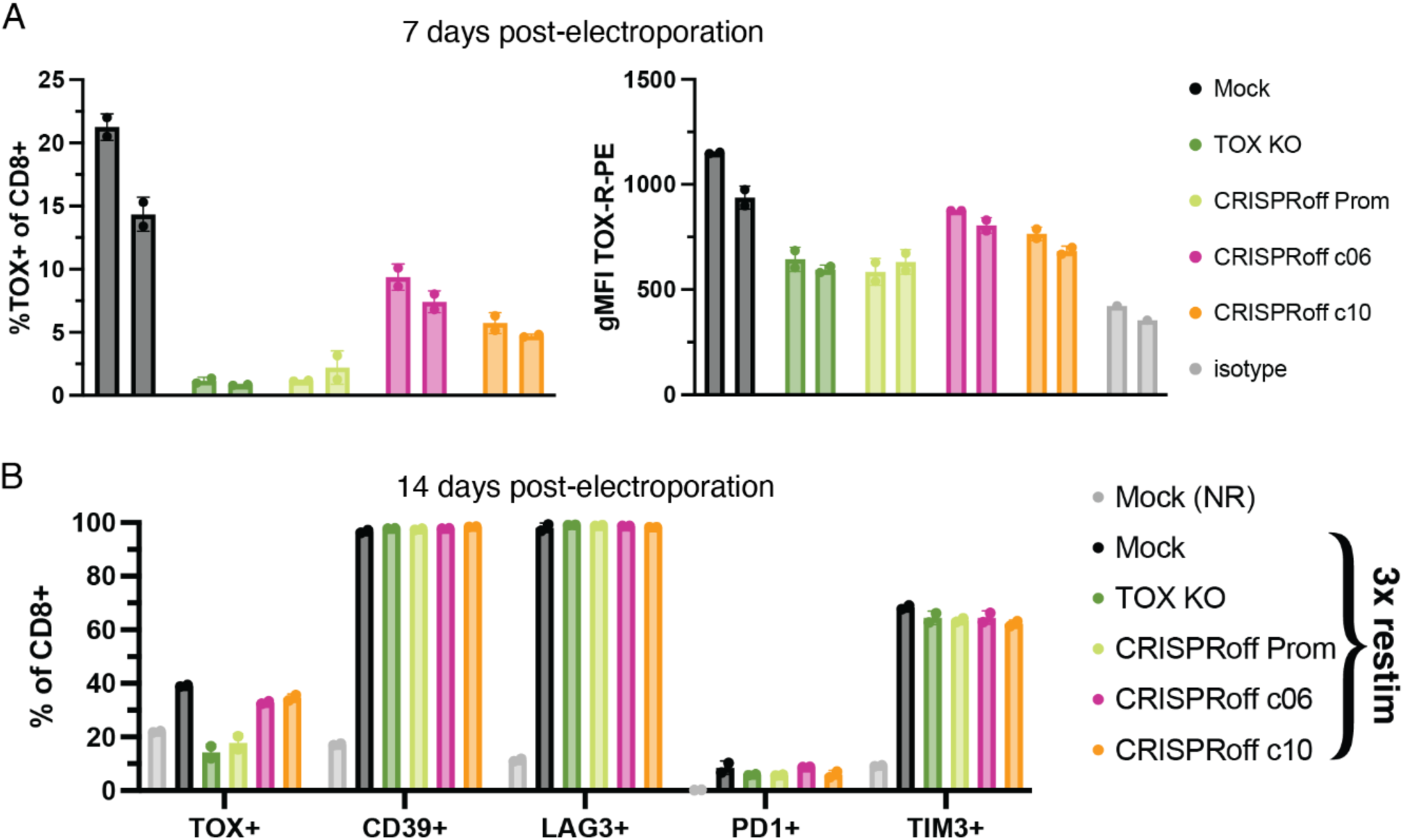
TOX modulation does not impact inhibitory receptor expression in CD8+ T cells repetitively stimulated with anti-CD3/anti-CD28 *in vitro*. A) TOX expression as quantified by flow cytometry 7 days post-electroporation. (Left) Percentage of CD8+ cells expressing TOX; (Right) geometric mean fluorescence intensity of TOX-R-PE. Each bar represents a distinct T cell donor (N=2) and the average of 2 electroporation replicates. B) Flow cytometry immunophenotyping of CD8+ T cells from N=1 T cell donor (compare to Fig. 3B) 14 days post-electroporation +/− 3x restimulation. Percentage of CD8+ cells expressing CD39, LAG3, PD1, TIM3, and TOX, respectively, were quantified. Bars represent the average of 2 electroporation replicates.

**Supplementary Figure 4.**
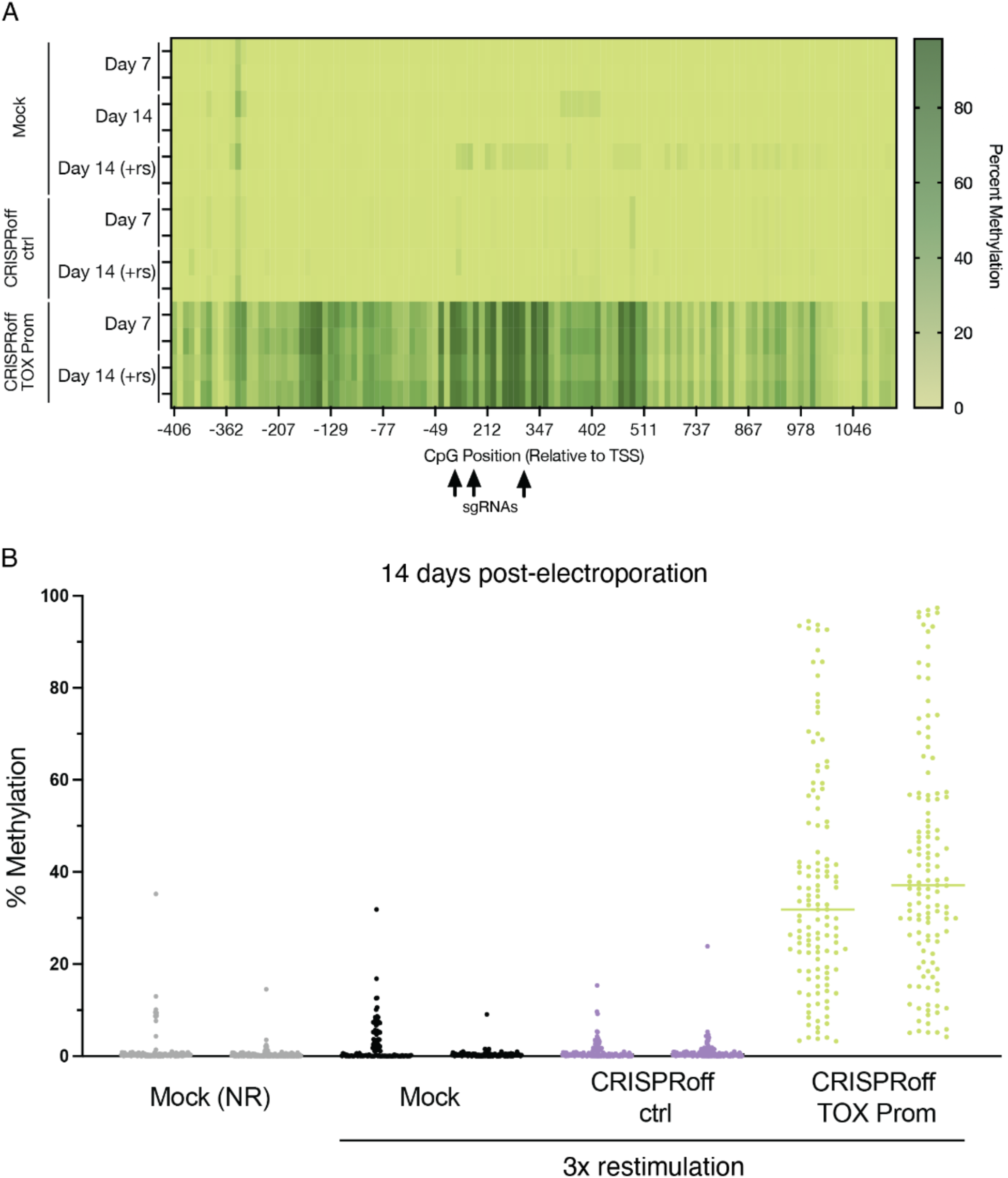
TOX promoter methylation is deposited after treatment with CRISPRoff and maintained after restimulation with anti-CD3/anti-CD28 *in vitro*. A) Heatmap depicting CpG methylation proximal to the *TOX* promoter before and after restimulation. N=2 T cell donors with 2 electroporation replicates per treatment condition. B) Average percent methylation per CpG dinucleotide in *TOX* amplicon as measured by EM-Seq 14 days after CRISPRoff treatment; N=2 T cell donors with 2 electroporation replicates per treatment condition.

**Supplementary Figure 5.**
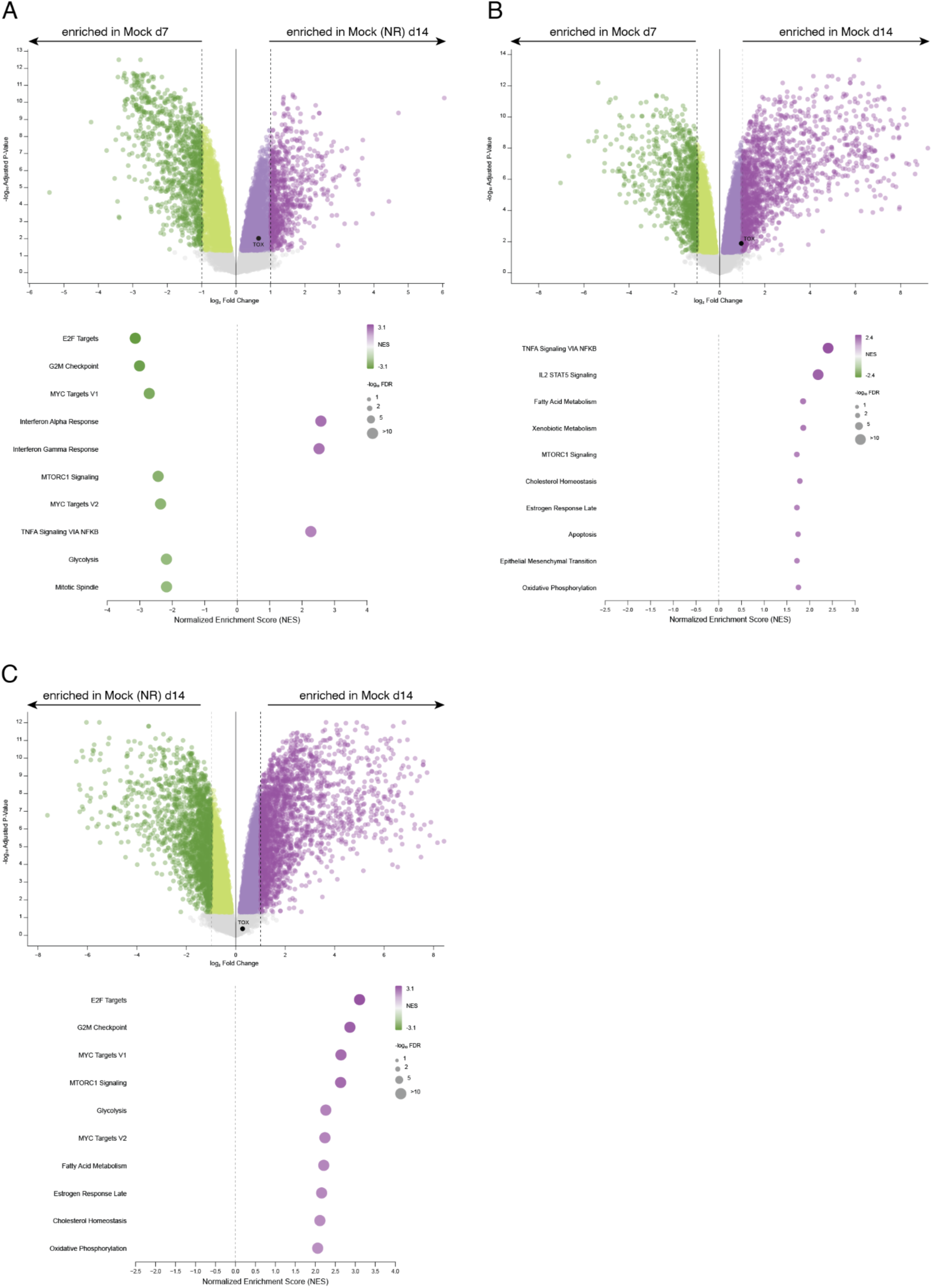
Repetitive stimulation with anti-CD3/anti-CD28 *in vitro* is associated with widespread transcriptional changes. A-C) (Top) RNA-Seq volcano plots depicting differential gene expression (FC>=2, adj. p-value<0.05) and (Bottom) RNA-Seq pathway enrichment for the following comparisons (N=3 T cell donors, 2 electroporation replicates per condition): A) Mock (NR) day 14 vs. Mock day 7; B) Mock day 14 vs. Mock day 7; C) Mock day 14 vs. Mock (NR) day 14.

## MATERIALS AND METHODS

### *In vitro* transcription of mRNA for electroporation

Templates for *in vitro* transcription were generated from plasmids by PCR amplification and subsequent purification. ABE8.20m nSpG plasmid (KAC1160) was provided by Ben Kleinstiver [22], primers and PCR amplification protocol were optimized by Jessica Nguy (NEB Phusion HotStart; F primer: GCCGCTAATACGACTCACTATAAGGGAGAGCCGCCACCATGGCCGCTAATACGACTCACTATAAGGGAGAGCCGCCACCATG; R primer: TTTTTTTTTTTTTTTTTTTTTTTTTTTTTTTTTTTTTTTTTTTTTTTTTTTTTTTTTTTTTTTTTTTTTTTTTTTTTTTTTTTTTTTTTTTTTTTTTTTTTTTTTTTTTTTTTTTTTTTAAGCCATAGAGCCCACCGCATCCCC; Tm= 70C; ext. time= 2 min). CRISPRoff-v2.3 plasmid [24], primers, and amplification protocol were provided by Laine Goudy (Alex Marson, Luke Gilbert). mRNAs were *in vitro* transcribed using HiScribe T7 mRNA kit with CleanCap Reagent AG (New England Biolabs #E2080) with N1-Methyl-Pseudouridine-5’-Triphosphate (TriLink, #N-1081) substitution according to mRNA synthesis protocol with modified nucleotides. After incubation with DNase I, mRNA was purified with MEGAclear Transcription Clean-up Kit (Invitrogen, #AM1908) according to kit instructions and stored at −80C.

### Single guide RNAs (sgRNAs)

spCas9 sgRNAs were ordered from IDT (with standard modifications) and resuspended in TE buffer to a concentration of 100 uM for use. See supplementary table 1 for spacer sequences.

Design and selection of adenine base editing guide: Adenine base editor guide RNAs were nominated using the base editor design tool (https://github.com/mhegde/base-editor-design-tool) and filtered for splice disrupting mutations by Jessica Nguy. A handful of guides were tested for efficacy (data not shown) and the highest efficiency guide (targeting the splice donor in intron 1) was selected for use here.

Selection of sgRNAs for CRISPRoff epi-silencing: “TOX Prom” pool consisted of TOX guides ranked 2-4 in the hCRISPRi-v2.1 library (supplementary table 3 in Horlbeck, *et al.* eLife 2016 [25]); the 5’ G was replaced with the endogenous nucleotide. For selection of guides targeting c01-c11 with CRISPRoff, enhancer predictions (cREs [27]) for TOX from 18 T cell datasets and 1 Jurkat data set (supp. table 2) were visualized as a custom track in the UCSC genome browser [49] along with CD8+ DNaseI hypersensitivity (DS17203 [50]). Guides were manually selected from the CRISPR targets track [51–55] to tile regions where DNaseI hypersensitive sites overlapped enhancer predictions. A minimum of 4 guides were selected per region.

### T cell sourcing

T cells were isolated fresh from healthy human donor PBMC leukopaks (StemCell) by washing with EasySep buffer (1X PBS without calcium or magnesium, 2% FBS, 1 mM EDTA) before negative selection using EasySep human T cell (StemCell, #17951), CD8+ (StemCell, #17953), or Naïve CD8+ (StemCell, #17968) isolation kit (according to manufacturer instructions).

### T cell activation and culture

Freshly isolated T cells were seeded at a density of 1×10^6^ cells/mL in T cell medium consisting of X-VIVO 15 (Lonza, #04-418Q), 5% FBS, 50 μM 2-mercaptoethanol, and 10 mM N-acetyl L-cysteine. T cell medium was supplemented with cytokines to a final concentration of 100 or 500 IU/mL of human recombinant IL-2 (PeproTech, #200-02) as indicated; with or without addition of 5 ng/mL human recombinant IL-7 (PeproTech, #200-07) and 5 ng/mL human recombinant IL-15 (PeproTech, #200-15) as indicated. CD3/CD28 Dynabeads (Gibco, #11132D; #40203D) were added to culture in a quantity equivalent to the total number of T cells. T cells were cultured in appropriate cell culture flasks and incubated at 37°C, 5% CO2, 90% humidity. After activation, T cells were passaged to approximately 5×10^5^ cells/mL in T cell medium supplemented with IL-2 every 2-3 days. T cell restimulations were performed with Immunocult (StemCell, #10970) or CD3/CD28 dynabeads (Gibco, #11132D; #40203D).

### T cell electroporation

2-3 days after activation, T cells were removed from beads using magnetic separation. After manual counting by trypan blue on a hemocytometer, an appropriate quantity of T cells (5-7.5×10^5^ per well) was centrifuged at 400xg for 5 minutes, washed once with 1X PBS (without calcium or magnesium), and then resuspended in freshly prepared P3 buffer (20 uL/well), mixed with editing reagents (1-2 ug editor mRNA + 50 pmol sgRNA), and transferred to 96-well nucleocuvette plate (Lonza, #V4SP-3960). T cells were electroporated using the pulse code DS137 on the core unit + 4D-Nucleofector 96-well unit. After nucleofection, warm T cell medium (without additives) was added to wells of the nucleocuvette plate and T cells were rested inside the incubator for 15 minutes before being transferred to appropriate cell culture vessels with additional warm T cell media and IL-2 (final concentration of 100-500 IU/mL, as indicated).

### Reverse Transcription quantitative Polymerase Chain Reaction

RNA was isolated from T cells (appx. 0.2-1×10^6^ cells) using RNeasy micro kit (Qiagen, #74004) with DNase incubation and elution in 20 uL of RNase-free water. RNA concentration was quantified by Nanodrop to normalize RNA input to reverse transcription reactions. cDNA was synthesized using SuperScript IV VILO Master Mix according to kit instructions. After synthesis, cDNA was diluted in nuclease-free water. Duplex TaqMan assays were used to quantify transcript abundance (Thermo Fisher FAM-MBG: #4331182 Assay ID (TOX): Hs01049519_m1; VIC-MBG-PL: #4448485 Assay ID (B2M): Hs99999907_m1). Briefly, TaqMan assays were combined with water and SsoAdvanced Universal Probes SuperMix (BioRad, #1725281) and distributed to wells of a white qPCR plate (BioRad, #MLP9651) and mixed with diluted cDNA. The duplex assay was optimized such that the final concentrations of the B2M VIC-MBG-PL and TOX FAM-MBG assays were 0.5X and 1X, respectively. Each sample was run in technical triplicate. Plates were sealed well with optically clear tape (BioRad, #MSB1001), quickly spun, and run on CFX96 Touch Real-Time PCR Detection System using 2-step amplification protocol specified by SsoAdvanced Universal Probes SuperMix manual, run for 40 cycles. Analysis of transcript abundance was performed by normalized relative quantification by standard curve.

### Flow Cytometry

#### Proliferation staining

T cells were centrifuged at 300xg for 7 min and washed in PBS. Cells were resuspended in 1 mL of staining solution (2.5 uM CFSE or CTV in PBS) per 10 ×10^6^ cells. After a 10 min incubation at room temperature protected from light, staining was quenched with 9-10 volumes of 2% BSA in PBS. Cells were washed twice more before proceeding to culture or surface staining.

#### Surface staining

T cells (appx. 1-4×10^5^) were transferred to 96w U-bottom plate (fisher cat#) and washed twice with EasySep buffer or PBS (400-500xg 4-5 min) before surface staining with viability dye and fluorochrome-conjugated antibodies. Samples were incubated on ice for 20-45 min then washed twice with EasySep buffer. Surface-stained cells were resuspended in EasySep buffer for acquisition on the Attune Nxt cytometer or fixed for intracellular staining.

#### Intracellular staining (ICFC)

For intracellular staining with FOXP3/Transcription Factor Buffer Set (eBioscience #00-5523-00), surface-stained cells were resuspended in fixative and incubated on ice for 30 min. Following fixation, cells were washed twice with permeabilization buffer, then incubated for 15 min on ice in a blocking solution consisting of permeabilization buffer, normal rat serum (1:50, eBioscience #24-5555-93), and human TruStain FcX (1:20, Biolegend #422302). After blocking, intracellular stains were applied and incubated for 45 min on ice. After 2 more permeabilization washes, cells were resuspended in EasySep buffer for acquisition on the Attune Nxt cytometer.

See supplementary table 3 for antibodies/dyes used.

### Next Generation Sequencing

Genomic DNA was harvested from cells using DNeasy Blood and Tissue (0.1-1×10^6^ cells; Qiagen #69504) or QuickExtract (50 uL per 5×10^4^ cells; BioResearch Technologies #76081-768). Target regions were amplified by PCR with Q5 HiFi polymerase (New England Biolabs #M0492L) with primers containing barcoding adaptors, purified, and amplified again for barcoding to generate sequencing libraries. Library preparation was performed in house or by the Innovative Genomics Institute Next Generation Sequencing Core. Libraries were sequenced at the core on the Illumina MiSeq instrument with 2×300 bp reads (Illumina #MS-102-3003).

### Quantification of base editing

Following NGS, adaptor trimming of .fastq files was performed with fastp prior to analysis with CRISPResso2 to quantify editing efficiency and allele frequency.

### Enzymatic Methyl Sequencing

We performed EM-Seq using methods adapted from LEMONmethyl-Seq (Christenson, et al. bioRxiv 2026). Briefly, genomic DNA was harvested from cells using DNeasy Blood and Tissue (0.2-1×10^6^ cells, Qiagen #69504). Genomic DNA was then digested by treatment with restriction enzymes Not1-HF, NcoI-HF, and XhoI (New England Biolabs #R3189S, #R3193S, and #R0146S) and subsequent clean up with QiaQuick PCR Purification Kit (Qiagen #28104). EM conversion was performed with NEBNext Enzymatic Methyl-seq v2 Conversion Module (New England Biolabs #E8020S) according to kit instructions; all clean up reactions were performed with Ampure XP beads (Beckman Coulter #A63881). EM-converted DNA was then PCR amplified with KeyPo SE polymerase (Tm=58C, Ext 45s, 35 cycles; Vazyme #PK510-01) using EM-specific primers designed to target 1.5 kb overlapping the *TOX* TSS (designed using https://www.methprimer.com/). Amplicons were bead-purified and submitted for long-read sequencing at the UC Berkeley DNA Sequencing Facility by Oxford Nanopore Sequencing. Data was analyzed by LEMONmethyl-Seq computational analysis methods to quantify methylated cytosines.

### RNA Sequencing

Approximately 5×10^5^ cells were lysed in DNA/RNA shield and stored at −20C before sending to Plasmidsaurus for library preparation and RNA-Seq using 3’ end counting to quantify transcriptome-wide differential gene expression. Plasmidsaurus bioinformatics pipeline and user interface were used for data analysis and visualization.

### Data Availability

All data files, raw and processed, available upon request.

